# Proteomic profile of hippocampal growth cones through early postnatal development

**DOI:** 10.1101/2025.11.10.687591

**Authors:** Maike Krause, Kamilla Aase Kronberg, Paulo J. B. Girão, Lara Sophie Strohmeier, Giulia Quattrocolo

## Abstract

The early postnatal period represents a critical window for neural circuit assembly in the developing hippocampus. Growth cones (GCs) play essential roles in establishing connectivity patterns, yet their proteomic composition during this dynamic developmental phase remains poorly characterized. Here, we performed comprehensive proteomic profiling of GCs isolated from the whole hippocampus (HP-GCs) and dentate gyrus (DG-GCs) across the first five postnatal days (P1, P3, P5) in mice. Mass spectrometry and proteomic analysis revealed that GCs from both regions are highly similar at P1, subsequently diverging. This earliest timepoint seems to be characterized by a high metabolic demand. Developmental trajectories of the GCs from the two subregions then start to differ, with P3 being a very dynamic period for HP-GCs, with GO terms related to RNA splicing, and a transition towards synapse formation by P5. On the contrary, DG-GCs seem to still be in an exploratory phase by P5, suggesting a delayed maturation and reflecting the later time course of the invasion of EC layer 2 Reelin-positive axons into this region. We identified over 5,000 proteins, with each timepoint characterized by distinct protein subsets. Temporal trajectory analysis revealed key GC markers including DCC, DPSYL2, and FABP7, which showed dynamic regulation across development. Notably, we identified multiple proteins containing nuclear localization sequences, suggesting bidirectional communication between GCs and the nucleus during critical developmental periods. Furthermore, our dataset contains over 400 proteins associated with neurological and neurodegenerative diseases, with twenty showing temporal regulation, highlighting the potential relevance of early developmental disruptions to disease pathogenesis. These findings provide a comprehensive proteomic resource for understanding GC function during hippocampal circuit formation and offer insights into the molecular mechanisms underlying region-specific developmental timelines in the brain.

## 2. Introduction

The formation of precise neural connections during development depends critically on the ability of growing axons to navigate complex environments and establish appropriate synaptic contacts with their target cells. This process is mediated by growth cones (GCs), highly dynamic structures located at the tips of extending axons and serving as both sensory organs and motile organelles (Goodman & Shatz, 1993; Ramón y Cajal, 1890). GCs integrate multiple guidance cues from their environment, including attractive and repulsive signals, to direct axon pathfinding toward appropriate targets while avoiding inappropriate connections. The molecular machinery involved in the GC movement are complex signaling cascades that link extracellular guidance cues to cytoskeletal rearrangements (Dickson, 2002; Govek et al., 2005; Luo, 2000; Pérez-Ferrer & Herrera, 2025; Soriano et al., 2021; Tessier-Lavigne & Goodman, 1996). GC responses to guidance cues are highly context-dependent and can be modulated by the developmental stage of the neuron, the local molecular environment, and the activity state of the GC itself (Koppers & Holt, 2022). This adaptability is particularly important during the early postnatal period when circuit connection are established (Katz & Shatz, 1996; Zhang & Poo, 2001).

All these complex and dynamic processes that the GCs undergo are to be reflected in the proteins locally expressed. *In vitro* and *in vivo* studies have highlighted the ability of growth cones to quickly respond to changes in the encountered cues, regulating both their transcriptomic and their proteomic content (Cagnetta et al., 2018; Koppers et al., 2019; Pérez-Ferrer & Herrera, 2025; Shigeoka et al., 2016). However, thorough developmental analyses of the growth cone proteome *in vivo* are limited (Cagnetta et al., 2018; Chauhan et al., 2020; Dumrongprechachan et al., 2022; Estrada-Bernal et al., 2012; Nozumi et al., 2009; Poulopoulos et al., 2019). In this study we decided to focus on hippocampal growth cones, analyzing the changes in proteomic content within a short but critical developmental window.

The hippocampal circuit represents one of the most extensively studied neural networks in the mammalian brain, serving as a fundamental substrate for spatial navigation, episodic memory formation, and temporal coding (Moser et al., 2008). This circuit comprises a complex network of interconnected structures, including the entorhinal cortex (EC) and hippocampus (HP) with hippocampal subfields CA1 to CA3 and the dentate gyrus (DG) (Amaral & Witter, 1989).

The development of entorhinal-hippocampal connectivity follows a precise temporal sequence, with the establishment of major projection pathways occurring during embryonic and early postnatal periods in rodents (Amaral & Dent, 1981; Del Río et al., 1997; Fricke & Cowan, 1977; Supèr & Soriano, 1994; Witter et al., 2017). Reelin-positive Layer 2 neurons of the EC project primarily to the outer part of the molecular layer (ML) of the DG and to the stratum lacunosum-moleculare (SLM) of CA3, while layer 3 neurons predominantly target CA1 and the subicular complex (Steward & Scoville, 1976; Witter et al., 1988). In parallel with the establishment of entorhinal inputs, the connectivity within the HP itself develops. Dentate granule cells start to extend their axons (mossy fibers) toward their CA3 target soon after birth. These axons traverse the hilus and establish synaptic contacts with CA3 pyramidal neurons while avoiding inappropriate targets (Amaral & Dent, 1981; Blackstad et al., 1970). Between P2-P5, CA3 pyramidal cells start to form their connection with CA1 pyramidal cells by extending their fibers, forming the Schaffer collaterals. This laminar specificity is established during early postnatal development through precise axon guidance mechanisms and activity-dependent refinement processes (Canto et al., 2008; Förster et al., 2006; Mata et al., 2018).

Invasion of the HP by its major input cells requires precise coordination of attractive and repulsive guidance signals, with molecules such as Reelin, slit proteins, and semaphorins playing critical roles in directing axon pathfinding (Del Río et al., 1997; Förster et al., 2002; Mata et al., 2018). How these changes are dynamically represented in hippocampal GCs is unknown. To target this issue, we aimed to characterize GCs from the whole HP and the DG at a proteomic level. As the input of layer 2 EC is the major input to the DG and CA3 regions, we first characterized the timeline of ingrowth of those fibers by using a recently developed cell-type specific mouse line (Blankvoort et al., 2018). This analysis helped us to narrow down three critical timepoints for our study (P1, P3, and P5) that we selected to isolate GCs from the HP and the DG of wild type mice. Subsequent mass spectrometry and proteomic analysis revealed specific developmental trajectories for the different subregions and highlighted distinct functional states of HP-GCs and DG-GCs.

## 3. Results

### 1. Fibers of Reelin-positive layer 2 cells invade their target structures in the hippocampus during the first postnatal days

Connections among the different subregions of the hippocampal formation are established and refined in the first few weeks of postnatal development (Donato et al., 2017; Dumas, 2005). One critical event in this process is the invasion of the hippocampus (HP) and its subregions by its major input, the Reelin-positive layer 2 cells of the entorhinal cortex (EC). An early study identified postnatal day (P) 3 to P6 as critical window for the invasion of rat Dentate Gyrus (DG) by utilizing radioactive dyes. However, using an elegant approach to entangle this critical developmental window, this study was limited in its timeline and precision in labeling the invading fibers (Fricke & Cowan, 1977). To overcome those limitations and to determine the dynamic of the ingrowth of Reelin-positive layer 2 fibers, we took advantage of a recently developed transgenic mouse line which conditionally expresses the tetracycline-controlled activator protein (Tta) in Odz3-expressing Reelin-positive cells of entorhinal layer 2 (Blankvoort et al., 2018). To label the axons of interest, we crossed this line with a TetO-GCamp6 mouse line (Blankvoort et al., 2018) and performed immunohistochemical staining for GCamP6 at P1, P3, and P5. A control staining for Reelin and GCamP6 expression in this cross can be found in SFig. 1A. Our analysis showed that input fibers are present in the Cornu Ammonis (CA) regions of the HP already at P1, with the first fibers already reaching the CA3 subregions. At P3, we observed fibers in the inner blade of the DG, starting to invade the outer blade at P5 (Figure 1A). The increase in numbers of invading fibers at the analyzed timepoints corresponded to an intensification of the fluorescent signal in the different hippocampal subregions (Figure 1B). However, to further confirm that the increase in fluorescent signal was indeed related to new axons invading the specific subregions, we performed immunohistochemical labeling for Syntaxin7 (Stx7), a well-known GC marker (Nozumi et al., 2009). Colocalization of GCamP6 and Stx7 signals allowed us to identify GCs of axons from EC Reelin-positive layer 2 fibers (Figure 1C). In total, we counted 1734 GCs for P1, 1970 GCs for P3, and 8256 GCs for P5 showing that within the early postnatal days the number of EC fibers arriving in the HP increases significantly (SFig. 1C). Indeed, at P1, the majority of GCs in the target regions were found in the CA3 SLM, close to border with the inner blade of the DG. At the subsequent stages, the fraction of GC increased in the inner (P3) and outer blade (P5) of the DG (Figure 1D).

**Figure 1:**
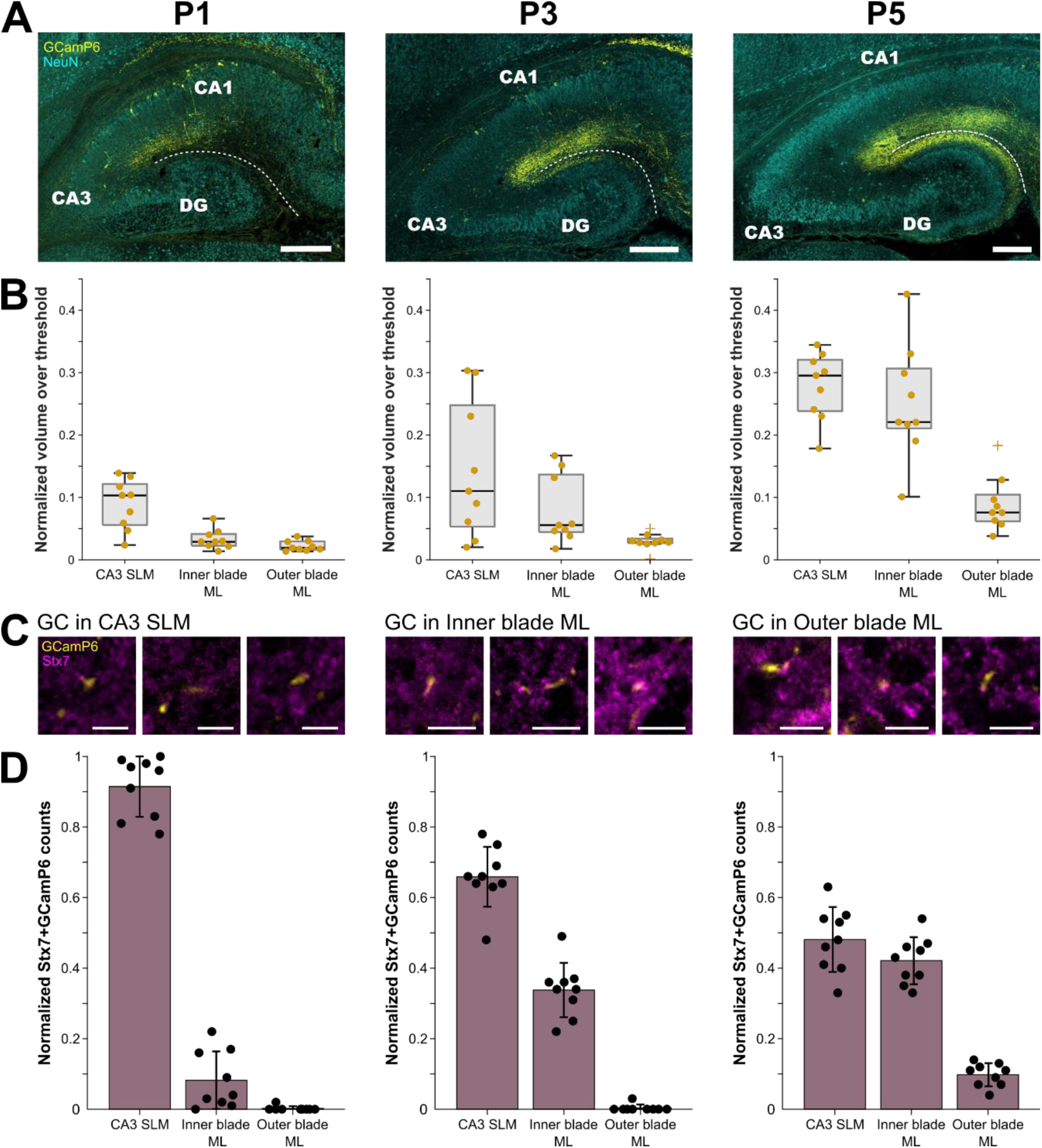
Invasion of the HP by fibers of entorhinal Reelin-positive layer 2 cells. A) Confocal image of horizontal sections of the HP of Odz3-tta;TetO-GCamP6 animals at P1, P3, P5. NeuN staining depicted in turquoise is used to label neuronal cell bodies. GCamP6 staining labels fibers of Reelin-positive layer 2 cells, shown in yellow. Scalebar: 200 µm. B) Quantification of the increase in GCamP6 fluorescent signal in CA3 SLM, and the molecular layer (ML) of the inner and outer blade of the DG over time reflecting how the invasion progresses during the early postnatal days. Datapoints identified as outliers are depicted as crosses. C) Representative confocal images of GCs of Reelin-positive layer 2 cells in CA3 SLM at P1 (left), inner ML at P3 (middle), and outer ML at P5 (right). GCs are indicated by Stx7 puncta (magenta) at the end of GCamP6 positive fibers (yellow). Scalebar: 5 µm. D) Quantification of layer 2 GCs density. Three nonconsecutive slices of three different animals were counted for the Stx7+GCamP6 events for each timepoint (n = 9). Given numbers are normalized to total counts per slice.

These findings demonstrate a clear sequential invasion of HP subregions by axons of the EC Reelin-positive layer 2 cells with the DG being the final target, with P3 to P5 being a potential particularly critical timepoint for the development of the connectivity in the DG. We therefore decided to use these three timepoints for our proteomic GC analysis.

### 2. Quality control experiments for Growth cone extraction of microdissected HP and DG

To study the GCs during early postnatal development of the HP on a proteomic level, we aimed to extract GCs from the total HP and the DG of wild type mice to submit them to mass spectrometry (Figure 2A). Before this crucial step, we performed a series of control experiments to assess different steps of our isolation protocol.

**Figure 2:**
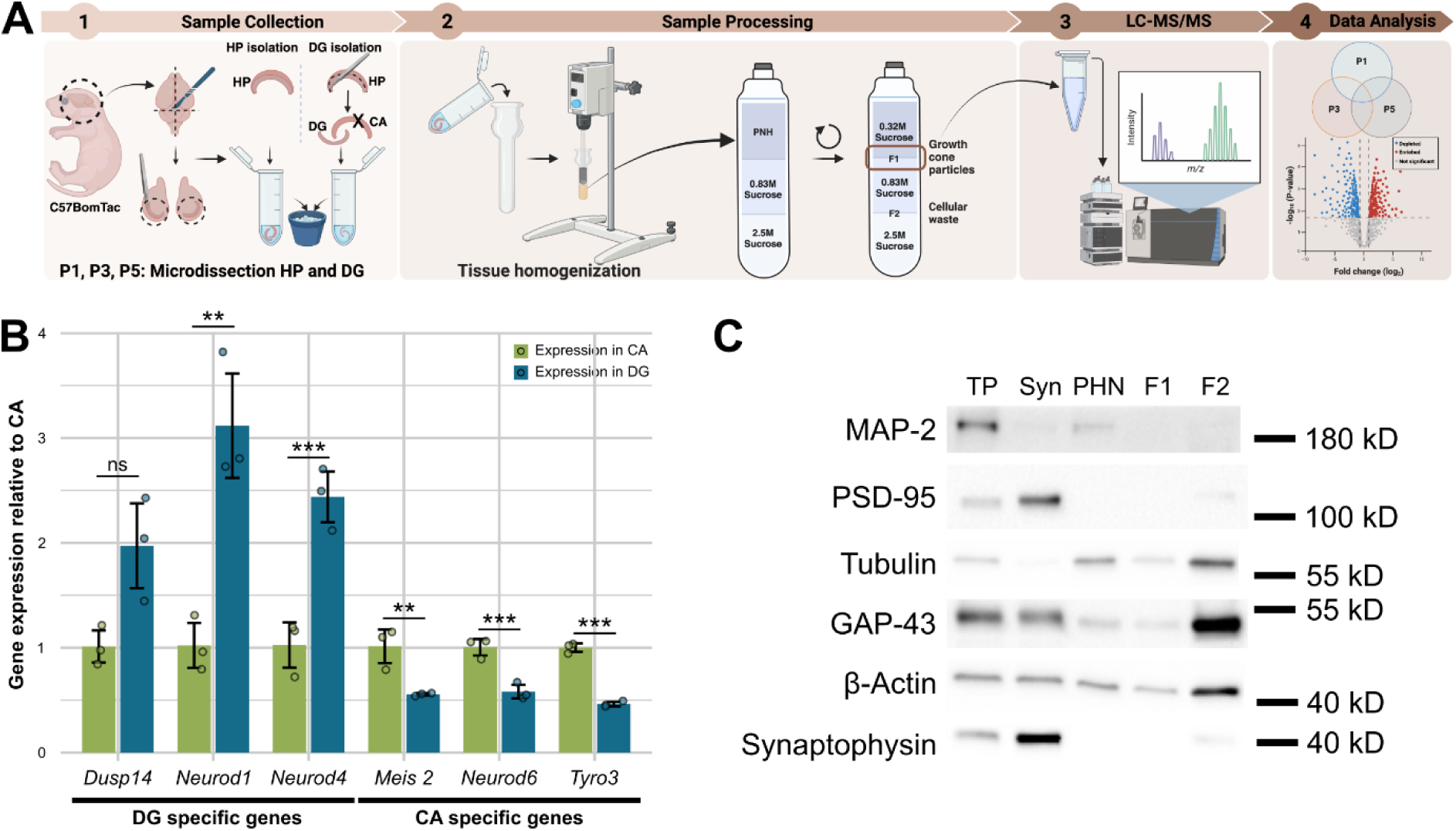
Validation of growth cone isolation protocol A) Overview for the sample preparation workflow (generated with BioRender). B) qRT-PCR results to verify the precision of DG dissection. Depicted is the relative gene expression to the CA tissue for DG-associated genes (*Neurod1*, *Neurod4*, and *Dusp14*) and CA-associated genes (*Meis2*, *Neurod6*, and *Tyro3*). Significance was tested with an independent two-tailed Student’s t-test and significance levels are indicated as *p<0.05, **p<0.01, and ***p<0.005, ns is non-significant. C) Western blot analysis as quality control for growth cone preparation. Post nuclear homogenate (PNH), F1, and F2 fraction after density gradient centrifugation are analyzed in comparison to the total hippocampal protein extract (TP) and adult synaptosome preparations (Syn). 1 µg of protein was loaded per sample. Detected were the neuronal marker MAP-2, the pre- and postsynaptic markers Synaptophysin and PSD-95, the cytoskeleton components Tubulin and β-Actin, as well as the GC marker GAP-43.

First, proper tissue dissection was verified with qRT-PCR using genes that are specific to the CA or DG region of the HP at P4. Genes were picked based on in situ hybridization data of the Allen developing mouse brain atlas (https://developingmouse.brain-map.org/). The dual specificity phosphatase 14 (*Dusp14*) was used as a marker for DG tissue, alongside three members of the Neurod family, a transcription factor family that is critical in tissue development and maintenance (Tutukova et al., 2021): *Neurod1* (Neuronal differentiation 1), and *Neurod4*, showed specific expression for the DG and *Neurod6* for the CA region. Further CA markers were *Meis2*, a homeodomain transcription factor, and *Tyro3* (Tyro protein tyrosine kinase 3), which are well known adult CA markers (Hagihara et al., 2009). DG-associated genes showed a significantly higher mRNA level for DG tissue while CA-associated genes were clearly more abundant in the CA tissue extracts, confirming our ability to correctly dissect the DG from the HP even at these early developmental stages (Figure 2B).

GCs from total HP and the DG were purified following a modified protocol by Pfenninger et al. (1983). After sucrose density gradient separation, GCs were harvested from the F1 fraction, while remaining components of a nuclear cleared tissue homogenate were collected in the F2 fraction of the gradient. To validate this step of our protocol, we performed a Western blot analysis (Figure 2B) of total HP-GC preparation at P4. As controls adult hippocampal total protein extract (TP) and adult synaptosome preparation (Syn) were used. As expected, PSD-95 and Synaptophysin were only detectable in adult controls. MAP-2 was majorly detected in mature tissue and only faintly represented in the Post nuclear homogenate (PNH), however, MAP-2 could not be detectable in any other GC fraction. Tubulin and β-Actin as important components of the cytoskeleton can be detected in almost all samples, except for the synaptosome sample that did not show any Tubulin signal. GAP-43 as classical growth cone marker can be detected in all fractions. Summarizing, we detect all GC associated proteins (GAP-43, Tubulin, and β-Actin) in the F1 fraction and none of the other makers. Collectively, our control experiments confirmed the extraction of high-quality GC of microdissected DG and HP (Figure 2).

### 3. Hippocampal growth cones in early postnatal days

To study how the proteomic content of hippocampal growth cones changes during early postnatal development, we generated five samples for hippocampal GC (HP-GC) at P1, P3, and P5, pooling both HP of a single animal for subsequent GC extraction. Extracted GC were filtered on 0.1 µm filters and submitted to LC-MS/MS for mass spectrometry analysis. Samples were measured in two separate batches of three and two biological replicates each. LC-MS/MS raw data was interpreted using MaxQuant v.2.6.1 (Cox & Mann, 2008) and subsequent data analyses were conducted with log2 transformed LFQ intensities with R. Proteins were counted as identified when 3 out of 5 replicates contained an LFQ intensity. Comparison of the log2 LFQ intensities for known GC markers in our dataset between the two batches showed no differences, thus, we considered no batch effect in the dataset.

For HP-GCs we identified 4,894 proteins in total with an average of 3,500 to 4,000 proteins per timepoint (Figure 3A). Venn diagram analysis highlighted subsets of protein that are timepoint specific (Figure 3B). At P1, we detected the largest subset of time specific proteins, with almost 800 proteins, while at P5 we identified only approximately 100 proteins. Comparing our data with a comprehensive GC marker list (Chauhan et al., 2020) revealed that approximately 80% of known GC markers (GCM) are represented in our dataset, a value similar to what was previously reported in the literature (Chauhan et al., 2020). We further assessed the reliability of our GC dataset by analyzing the KEGG pathways (FDR 0.05) covered by the proteome shared by the three different timepoints (Figure 3C). By focusing on the Top20 KEGG pathways (disease terms excluded), we identified several pathways expected to be associated with GCs, such as *Regulation of actin cytoskeleton*, *Focal adhesion*, *Axon guidance*, as well as *Ras* and *Rap1 signaling pathways.* Interestingly, we also identified *Ribosomes* and -*Spliceosomes* among the most enriched and most significant pathways in the shared proteome. These molecular machineries are critical for local mRNA modification and translation, enabling the GCs to rapidly remodel their local proteome in response to dynamic guiding cues (Cagnetta et al., 2018; Shigeoka et al., 2016).

**Figure 3:**
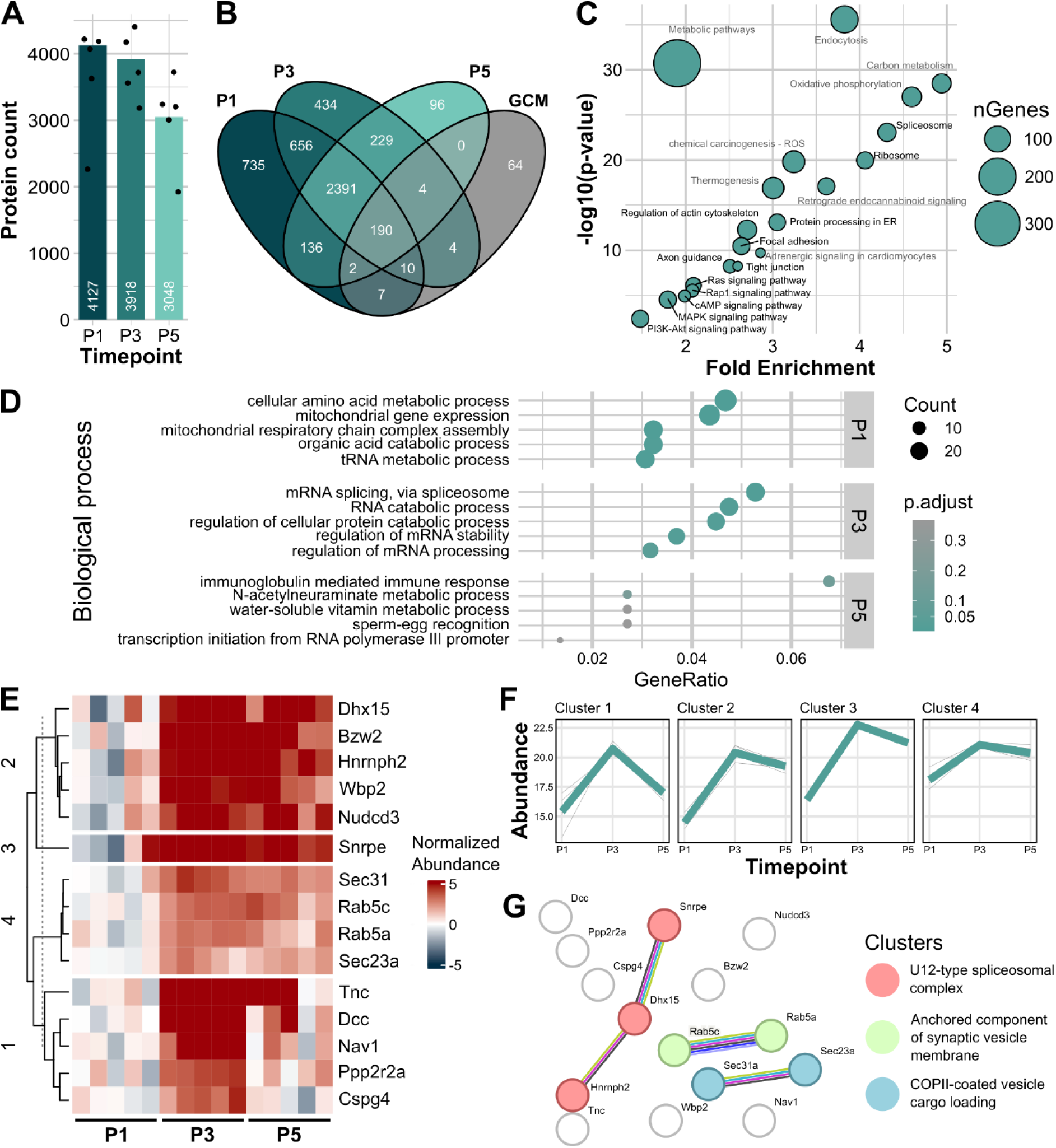
Proteome of HP-GC. A) Bar graph representing the number of identified proteins per timepoint (n = 5 animals for each timepoint). B) Venn diagram showing the distribution and overlap of identified proteins for each timepoints, and the overlap with a list of GC markers (GCM) from Chauhan et al., 2020. C) KEGG pathways identified for the proteome shared across timepoints. Relevant GC pathway terms are highlighted in black. D) List of Top5 Gene Ontology (GO) terms for biological processes for the timepoint specific proteins at P1, P3, and P5. Dot size indicates the protein count per term and the dot color the adjusted p-value. E) Heatmap representing expression across development normalized to the mean log2 LFQ intensity (abundance) at P1. Clusters were defined using the single time series regression analysis and k-means clustering. See also STab. 1. F) Detailed temporal trajectories for each cluster for the single time series temporal trajectory analysis. Average Log2 abundance for all proteins in the cluster represented in teal and the individual protein signals in grey. G) STRING network analysis of all significant proteins of the temporal analyses with all known interactions between proteins depicted in default color code. Three functional clusters were identified within the STRING network.

To understand which biological processes our identified proteins are related to, we performed a Gene ontology (GO) term analysis for the timepoint specific proteins for HP-GCs at P1, P3, and P5 (Figure 3D). At P1, the most abundant terms were associated with mitochondrial processes such as *Mitochondrial gene expression* and *Mitochondrial* r*espiratory chain complex assembly*. GCs have a high energy demand and need a substantial amount of ATP for active axon extension, cytoskeletal remodeling, and membrane synthesis. The enrichment of *Mitochondrial respiratory chain complex assembly* could suggest that a robust energy-producing machinery is present to cover the high energy demands (reviewed in: Smith & Gallo, 2018). In addition, *Mitochondrial gene expression* is indicative for the production of mitochondria to meet the metabolic needs of GCs (Vaarmann et al., 2016). Interestingly, *tRNA metabolic processes* were also highly enriched, underlying the high demand for efficient protein synthesis at early timepoints (Estrada-Bernal et al., 2012).

The GO term analysis for P3 specific proteins revealed biological processes mainly associated with *RNA catabolic processes* and *RNA splicing*. Earlier studies on GCs showed that the active response to guiding cues and the establishment of synaptic contacts requires a rapid turnover of specific mRNAs (Shigeoka et al., 2016). Alternating available mRNAs via Splicing would allow a GC to fine tune its response by generating developmentally relevant isoforms of key proteins, such as cell adhesion molecules and guidance receptors (Z. Chen et al., 2008). This would not only contribute to circuit formation (Shigeoka et al., 2016) but could also indicate molecular reprogramming where most HP-GCs enter a new developmental phase, transitioning from pathfinding and dynamic exploration to stable connectivity. While the biological processes identified for the P5 specific terms showed overall a lower degree of significance, the first term, *Immunoglobulin mediated immune response,* stood out. This term is triggered among others by the protein C1qb, the core component of the complement C1 complex. Interestingly, this protein complex is not only important for the immune response itself, but also crucial for microglia-mediated synaptic pruning, a process with immune-like mechanisms that eliminate excess synaptic connections in GCs and developing circuits (Stevens et al., 2007). The appearance of these specific terms only at P5 could further suggest a functional transition of HP-GCs from the exploration phase to the formation of stable connections.

To better evaluate the dynamicity and temporal changes in the HP-GC proteome we conducted a single-time series analysis utilizing the maSigPro R package (Conesa & Nueda, 2025). All proteins were included in the analysis, setting the minimum observations to four, with P1 used as reference timepoint. After a least square linear regression fit of the log2 LFQ intensities (abundance) and a Benjamini-Hochberg adjusted p-value below 0.05, fifteen proteins were identified with significant changes between P1, P3, and P5 in all biological replicates (Figure 3E). Using k-means clustering, we were able to best represent the individual changes with four clusters. All clusters showed a similar pattern of increase from P1 to P3, followed by a more or less steep decrease from P3 to P5. Indeed, almost all proteins identified in the single-time series analysis showed peak abundance around P3 indicating this timepoint as critical in the maturation of HP-GCs.

To highlight possible interactions, we performed STRING network analysis for all fifteen proteins together, obtaining three functional networks (Figure 3F). Of these, one included the *U12-type spliceosome complex*, a complex processing a small subset of introns in genes often involved in cell cycle regulation and development (Edery et al., 2011; Otake et al., 2002). Interestingly, this network also includes the heterogeneous nuclear ribonucleoprotein H2 (HNRNPH2), a protein known to have a Nuclear Localization Signal (Gonzalez et al., 2023), hinting at a mechanism of communication between the growth cones and the nucleus to regulate transcription. This protein is also known for its regulatory effect on alternative splicing (Rappe et al., 2014). The second network identified, *Anchored components of synaptic vesicle membrane,* seems again to highlight the transition to active synaptogenesis and synaptic maturation. Finally, *COPII-coated vesicle cargo loading* is associated with ER-to-Golgi transport, indicating increased protein trafficking to the GCs. Of special interest in this network is the known GC marker SEC31A (outer shell component of COPII) (Chauhan et al., 2020). Not included in any functional network but identified among the fifteen temporal changing proteins is DCC (Netrin receptor DCC), a protein whose role in axonal targeting has been well characterized and is known to mediate axon attraction upon ligand binding (Powell et al., 2008; Ren et al., 2007). Furthermore, DCC can bind to ribosomes and, thus, also regulate local translation in axons (Koppers et al., 2019).

Together, these observations indicate a coordinated temporal regulation of mitochondrial activity, RNA processing, and protein trafficking during the early postnatal days of the HP-GC development. The maSigPro analysis highlights P3 as a potential critical timepoint marking the transition from axonal growth and exploration to synaptic maturation.

### 4. Growth cones of the DG in early postnatal days

Knowing that the DG is the last subregion in the HP receiving its input from the layer 2 cells of the EC, we were interested in what the characteristics of the GC proteome in the DG are. As for the HP-GCs, we sampled wild type pups at P1, P3, and P5, pooling both DG of one animal as one biological replicate. Five biological replicates were generated for each timepoint and analyzed in two batches as previously described for HP-GCs. We identified approximately 4,000 proteins for the DG-GC samples from P1, P3, and P5 (Figure 4A), and a total of 4,987 proteins overall. Similarly to what we detected for the HP-GCs dataset, we found a significant number of proteins that were only identified for one of the three timepoints, again with P1 showing the highest number of time specific proteins. Interestingly the number of time specific proteins is lowest at P3 in DG-GCs, in contrast to what was observed for the HP-GCs, where the P5 sample showed by far the lowest number of time specific proteins. In fact, P5 in DG-GCs showed a very high number of time specific proteins, a number that was more than three times what we identified in HP-GCs. The Venn diagram for DG-GCs (Figure 4B) additionally revealed that the P3 and P5 samples showed a larger overlap in their proteome than the corresponding HP-GC samples (DG-GCs 414 proteins vs HP-DGs 239 proteins) suggesting that P3 and P5 DG-GCs might be more similar than the corresponding samples in HP-GCs. Also in this case, the comparison to the GC markers resulted in approximately 80% overlap, with similar results to what we saw for the HP-GCs. KEGG pathway analysis for the proteome shared by the three different timepoints revealed GC associated pathways and resulted in the same Top20 terms that were identified for the HP-GCs shared proteome, except for two terms *Chemokine signaling pathway* and *Adrenergic signaling in Cardiomyocytes* (SFig. 2).

**Figure 4:**
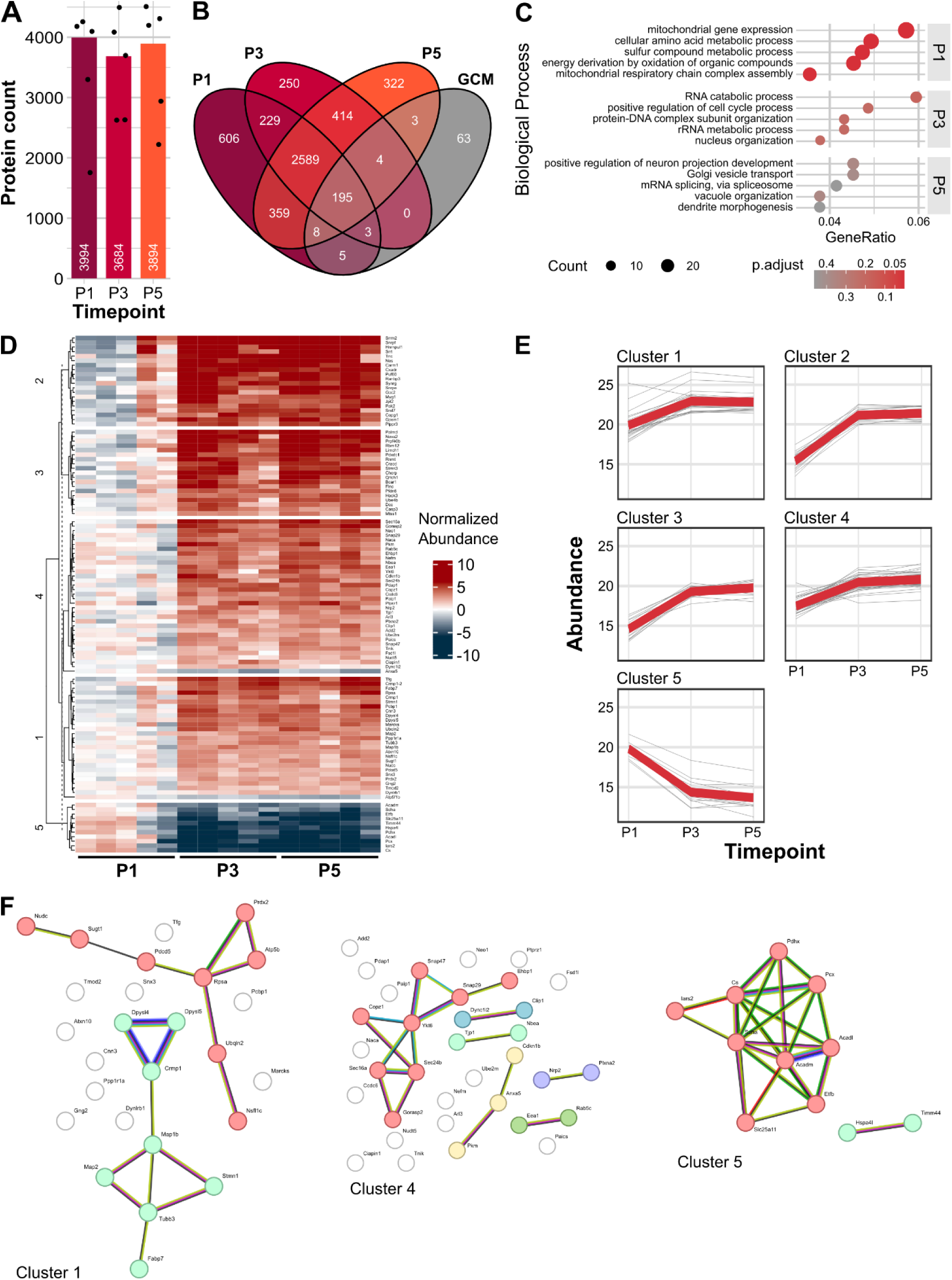
Proteome of DG-GCs. A) Bar graph representing the number of identified proteins per timepoint. B) Venn diagram showing the proteins identified for each timepoint and the overlap with the GC marker reference dataset of Chauhan et al. C) List of Top 10 results for the GO term analysis for timepoint specific proteins in the DG-GC dataset. D) Heatmap showing the normalized abundance and results of k-means clustering. 111 significantly changed proteins are represented. Protein names and their clusters can be found in STab. 1. E) Averaged (red curves) and protein individual (grey curves) changes in abundance for each single time series cluster. F) Results for functional network analysis for cluster 1, 4, and 5 conduced with STRING network analysis.

To achieve a deeper insight into the function of the identified proteins, we again ran a GO term analysis for the timepoint specific proteins of DG-GCs (P1: 606, P3: 250, and P5: 322) (Figure 4C). At P1, we observed similar terms to what previously discussed for the HP-GCs, with biological processes associated with *Mitochondrial gene expression* and *Respiratory chain complex assembly,* as well as tRNA metabolic processes also found for DG-GCs. This result might once again underscore that the GCs are in a state of high energy demand, active protein synthesis, and are in their exploration phase. Biological process GO terms identified for P3 and P5 specific proteins only showed minor significance. However, it is interesting to point out that the majority of terms identified for DG-GCs were different from the terms found for HP-GCs. At P3, among the highest rated terms are *RNA catabolic processes* and *Protein-DNA complex subunit organization*. Terms related to RNA metabolism are common between the two HP-GC and DG-GC P3 specific GO terms, suggesting P3 is very dynamic in what relates to RNA transcription. The term *Protein-DNA complex subunit organization* was triggered by members of the histone family (H2BC4) and their potential regulatory interaction partner (SET, ANP32E, ANP32B, TAF10). Histones have been observed in cerebellar neurites (Mishra et al., 2010) as well as in the axonal proteome of developing isolated retinal axons (Cagnetta et al., 2018; Moretti et al., 2015). The ANP32 protein family is known to cover a broad spectrum of functions, ranging from phosphatase inhibition and chromatin regulation to intracellular transport and it was shown that this family is also important for normal development in mice (Reilly et al., 2011). As previously mentioned, in the P3 HP-GCs, the major biological processes were related to local mRNA alteration via splicing. The splicing term was also detected for DG-GCs, however much less abundant (count) and less significant (adjusted p-value) and at a later timepoint (P5, Figure 4C). Besides, the analysis of GO terms of DG-GC at P5 highlighted two top terms, *Positive regulation of neuron projection development* and *Golgi vesicle transport*, suggesting that at P5 GCs from the DG are still in a process of dynamic exploration.

Summarizing, these observations on timepoint specific proteins reflect a shift in the timescale of GC development in the DG: while HP-GCs start to transition from mobile exploring structures to local entities forming first stable connection between P3 and P5, GC of the DG might still be in their exploration phase. This would reflect the timeline of invasion of the DG by its major input, the axons of entorhinal layer 2 cells, that have not fully invaded the outer blade of the DG until P5 (Figure 1A, P5).

To highlight the trajectory of DG-GCs, we ran a single time series analysis also for DG-GCs, which resulted in 111 significantly differently expressed proteins across the three analyzed timepoints (Figure 4D). The changes in abundance were best represented in five clusters utilizing k-means clustering (Figure 4E). While Cluster 1 to Cluster 4 exhibit increasing protein abundance over time, Cluster 5 is the only cluster that represents proteins that are decreasing in abundance at the later timepoints. The most abundant proteins overall can be found in Cluster 1. In fact, Cluster 2, Cluster 3, and Cluster 4 are primarily distinguished by their initial abundances at P1 and the following trajectory of increase.

To assess protein-protein interactions within each temporal profile, we performed STRING network analysis independently for each cluster (Figure 4F, SFig. 3). Two functional networks were identified in Cluster 1. One network (mint) is associated with both the *CRMPs in Sema3A signaling* and *Negative regulation of microtubule polymerization or depolymerization*. CRMPs are PlexinA-interacting proteins that act as mediator of SEMA3A signaling and neuronal differentiation (reviewed in Schmidt & Strittmatter, 2007). Phosphorylation of CRMPs inhibits their ability to bind tubulin dimers, leading to destabilization of the microtubule and ultimately causes the collapse of the GC (Arimura et al., 2005). Also part of this network is the Fatty acid binding protein-7 (FABP7), a brain specific lipid chaperone that plays essential roles during this developmental window by regulating the intracellular trafficking and metabolism of fatty acids and facilitating their transport to specific cellular compartments (Young et al., 2013). The second network (red) consists of the proteins ATP5B, NSFL1C, NUDC, PDCD5, PRDX2, RPSA, SUGT1, and UBQL2 and can be considered to be of mixed functionality. For example, NUDC (Nuclear migration protein nudC) plays a role in neurogenesis and neuronal migration (Aumais et al., 2001), while UBQL2 is known to be a key player in proteostasis. UBQL2 participates in several quality control pathways, such as the ubiquitin-proteasome system, ER-associated degradation, autophagy, and mitochondrial protein surveillance (UBQLN functions reviewed in Lin et al., 2022). ATP5B is part of the mitochondrial ATP synthases and contributes to energy metabolism (Chinopoulos et al., 2011), while RPSA is a small ribosomal subunit protein and can act as a surface receptor for laminin playing a role in cell adhesion and potentially being important for cell fate determination and tissue morphogenesis (Blazejewski et al., 2022).

Cluster 4 contains a total of six functional networks, with one major functional network (red) that represents the *SNARE complex* and *Snap receptor activity*. The SNARE (N-ethylmaleimide-sensitive factor attachment protein receptor) complex is mostly known for its involvement in membrane fusion and the synaptic vesicle cycle. Fusion of membrane vesicles managed by the SNAREs with coordinated elongation of the cytoskeleton is responsible for the growth of axons (reviewed in Ulloa et al., 2018). Another important STRING network in Cluster 4 represents the Semaphorin-plexin signaling pathway (violet) involved in neuronal protection guidance with the proteins Neuropilin 2 (NRP2) and Plexin A2 (PLXNA2). Both proteins are known to be crucial for the development of the hippocampal mossy fibers and DG (reviewed in Skutella & Nitsch, 2001; Zhao et al., 2018). The other networks covered by Cluster 4 are represented by few proteins, known to contribute to the retrograde microtubule transport (blue), establishment and maintenance of cell polarity, influence on the migration, proliferation of cortical neurons (mint and yellow), and cell adhesion and migration (green).

As mentioned above, Cluster 5 is the only cluster that showed a decrease in protein abundance in the single-time series analysis. As the majority of Cluster 5 proteins (red) are associated with *Fatty acid beta-oxidation using acyl-CoA dehydrogenase* and the *Citrate cycle*, this cluster represents key processes for energy production. In addition, the remaining two proteins in this cluster (mint), HSPA4L and TIMM44, are both involved in the import of proteins into mitochondria (Ting et al., 2017). Thus, Cluster 5 emphasizes a trend observed in the GO term analysis for the timepoint specific proteins, as at P1 protein synthesis and energy production were among the most important terms, disappearing by P3 and indicating a shift in functional needs of the GCs (Figure 4C).

Finally, Cluster 2 and Cluster3 contain one or two functional networks, respectively (SFig. 3). Cluster 2 network proteins are associated with *RNA processing* (red) and especially *mRNA splicing via spliceosome* (blue). Two STRING networks of Cluster 3 represent the pathway *Caspase activation via Dependence Receptors in the absence of ligand* and a second network without specific function (proteins: BCAR1 and FLNC). Remarkably of this cluster is that all proteins except of two are labeled as phosphoproteins. Among them are the earlier discussed DCC, the cytoskeleton interacting protein Stathmin-3 (STMN3, interacts with microtubules), Filamin C (FLNC, binds to actin), LIM and Calponin Homolog Domains 1 (LIMCH1, interacts with actin), Breast cancer anti-estrogen resistance protein 1 (BCAR1, general function in cytoskeleton remodeling), Metastasis suppressor 1 (MTSS1, promotes filopodia protrusions), and the protein degradation associated Ubiquitin Conjugation factor E4B (UBE4B), and Caspase-3 (CASP3). Modification of the phosphorylation status of those proteins would allow for a rapid and reversible response in protein localization (DCC), Protein-Protein interaction (BCAR1), cytoskeletal dynamics (STMN3, FLNC, LIMCH1, MTSS1), protein stability (FLNC, UBE4B), and enzymatic activity (CASP3, UBE4B), that could contribute the dynamic characteristics of GC during axon guidance and neuronal circuit formation (O’Donnell et al., 2009).

### 5. Subregion specificity of HP-GCs and DG-GCs

The analysis of proteome of GCs isolated from the whole HP and DG at early postnatal days showed that each timepoint has a specific subset of proteins that cover unique functions at P1, P3, and P5. P1 GCs of the HP and DG trigger similar functions in their GO term analysis, suggesting a high degree of similarity at this first stage. However, the GO term analysis for the time specific functions revealed for P3 and P5 that the timelines are shifted between the two different input tissues, indicating that overall, the HP-GCs develop on average earlier than the DG-GCs. Even the expression profiles of the different clusters analyzed of the time course analysis differs, with the HP-GCs clusters having a clear reduction in abundance from P3 to P5 (Figure 3F), in contrast to a much higher similarity in the DG-GCs clusters (Figure 4E). Therefore, to further understand the subregional differences in the two types of GCs we did a comparative analysis of the proteomes generated for HP-GCs and DG-GCs.

First, we analyzed the overlap between the timepoint specific proteins between the two datasets (Figure 3B and Figure 4B). The Upset plot (Figure 5A) shows the overlap between the six groups of timepoint specific proteins. The largest groups are the timepoint specific proteins at P1 for HP and DG. At P3, the number of timepoint specific proteins is decreased compared to P1 and at P5 further reduction was observed for the HP-GC but not for the DG-GC. Between HP and DG approximately 50% of the time-specific proteins at P1 are shared between the two input tissues, explaining the similar biological processes obtained in the GO term analysis for HP-GCs and DG-GCs time specific proteins. With increasing age, the overlap between P3 samples and P5 samples decreases from over 300 proteins to 50 and 7, respectively, further indicating that the HP-GCs and DG-GCs become less identical over time. Interestingly, the overlap between P1 HP-GCs with P3 and P5 DG-GCs is 30 and 18 proteins and the overlap between P3 HP-GCs with P5 DG-GCs is 75 proteins large (Figure 5A). This could support the presence of a different maturational trajectory of the DG compared to the HP and supports the observation that DG-GCs development is shifted to later timepoints compared to the HP-GCs (Figure 5B). In the group of proteins shifted to DG P5 we observed several members of the Transmembrane (TMEM) protein family. TMEMs are associated with various functions, such as maintaining cell homeostasis and cell-cell recognition (Q. Chen et al., 2021; Herrera-Quiterio & Encarnación-Guevara, 2023). TMEM106B, for example, is involved in protein degradation processes via lysosomes and a recent study demonstrated that a TMEM106B variant promotes neurite outgrowth and later spine formation in hippocampal neurons *in vitro* (Nguyen et al., 2023). Also TRIM46 was found at P5 in the DG and is crucial for the axon formation by regulation the microtubule organization (Harterink et al., 2019). This supports the idea that the GCs isolated at P5 from the DG are still in their elongating exploratory phase. Other interesting proteins found at later timepoints in the DG are PUM2, an RNA binding protein and master regulator of local translation (Hafner et al., 2010; Martínez et al., 2019), TMSB15B1 and TMSB15B2, both with assumed roles in cytoskeleton organization (Uniprot based on similarity), TOM1L1, that regulates endosomal trafficking, which is important for growth factor signaling (Wang et al., 2010), and COMMD1, associated with endocytic recycling and protein trafficking (Phillips-Krawczak et al., 2015). Since those proteins responsible for crucial functions in GCs are earlier detected in HP-GCs than DG-CCs this again emphasizes the temporal difference between the two GC populations we are studying. Interestingly, we also find a small number of proteins that are earlier detected in DG-GCs than HP-GCs, highlighting the fact that not only unidirectional processes from the CA subregions towards the DG are taking place during development in the HP, but also in the opposite direction (e.g. development of Mossy fibers).

**Figure 5:**
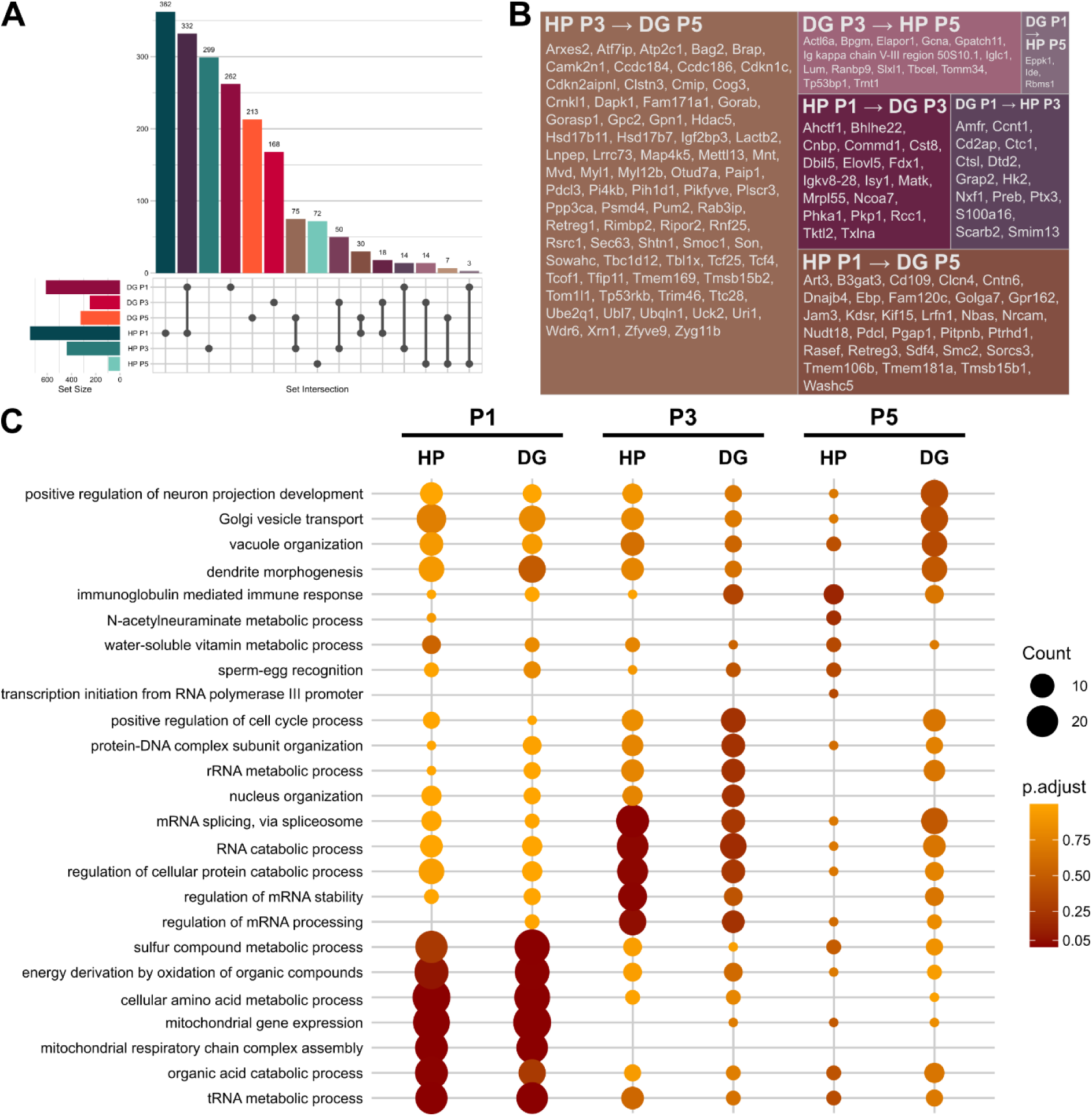
Comparative analysis of time specific proteins from HP-GCs and DG-GCs. A) Upset plot of the timepoint specific proteins for HP-GCs and DG-GCs. The lower left part of the plot represents the total number of proteins identified for P1, P3, and P5 in HP-GCs and DG-GCs. The lower right part represents the set intersection. Each set contributing to the intersection is marked with a point and connected with a line. The upperpart of the plot gives the number that belongs to the set intersection indicated below. B) Tile plot showing overlapping proteins between HP and DG at different timepoint. Color code based on A). C) Comparative depiction of the Top5 biological processes of HP-GCs and DG-GCs from Figure 3D and Figure 4C.

The temporal shift between the HP-GCs and DG-GCs is additionally highlighted when the Top5 biological processes for each timepoint are plotted together (Figure 5C). At P1 HP-GCs and DG-GCs share the same biological process. However, from P3 on, the Top5 terms drift apart. E.g., splicing associated proteins are still represented at P5 in the DG-GCs, while those terms almost disappeared for HP-GCs by this timepoint. At P5, HP-GCs are characterized by terms associated with target recognition while DG-GCs still show terms that are related to exploratory behavior.

Multi-time series analysis for DG-GCs against the HP-GCs enabled us to further identify tissue specific developmental changes in protein abundances. The analysis was conducted on a non-imputed dataset, as to avoid any possible assumption of the reason behind the missing proteins. Normalization to P1 HP-GCs values revealed abundances for 110 proteins significantly changing (p < 0.05) within the early postnatal days (Figure 6A). K-means clustering was best with eight clusters, with six clusters with increasing and two clusters with decreasing trajectories (Figure 6B). Cluster 1 and Cluster 8 represent proteins with the largest increase between P1 and P5. Smaller increases are represented in Cluster 2, Cluster 5, and Cluster 7 and moderately increasing changes are summarized in Cluster 3. Average expression for HP-GCs shows a peak abundance at P3 in Cluster 3, same is true for Cluster 1, Cluster 4, Cluster 7, and Cluster 8. No cluster shows a clear peak abundance at P3 for DG-GC proteins, somehow in line with what we previously observed (Figure 4E). Decreasing abundances were found in Cluster 4 and Cluster 6.

**Figure 6:**
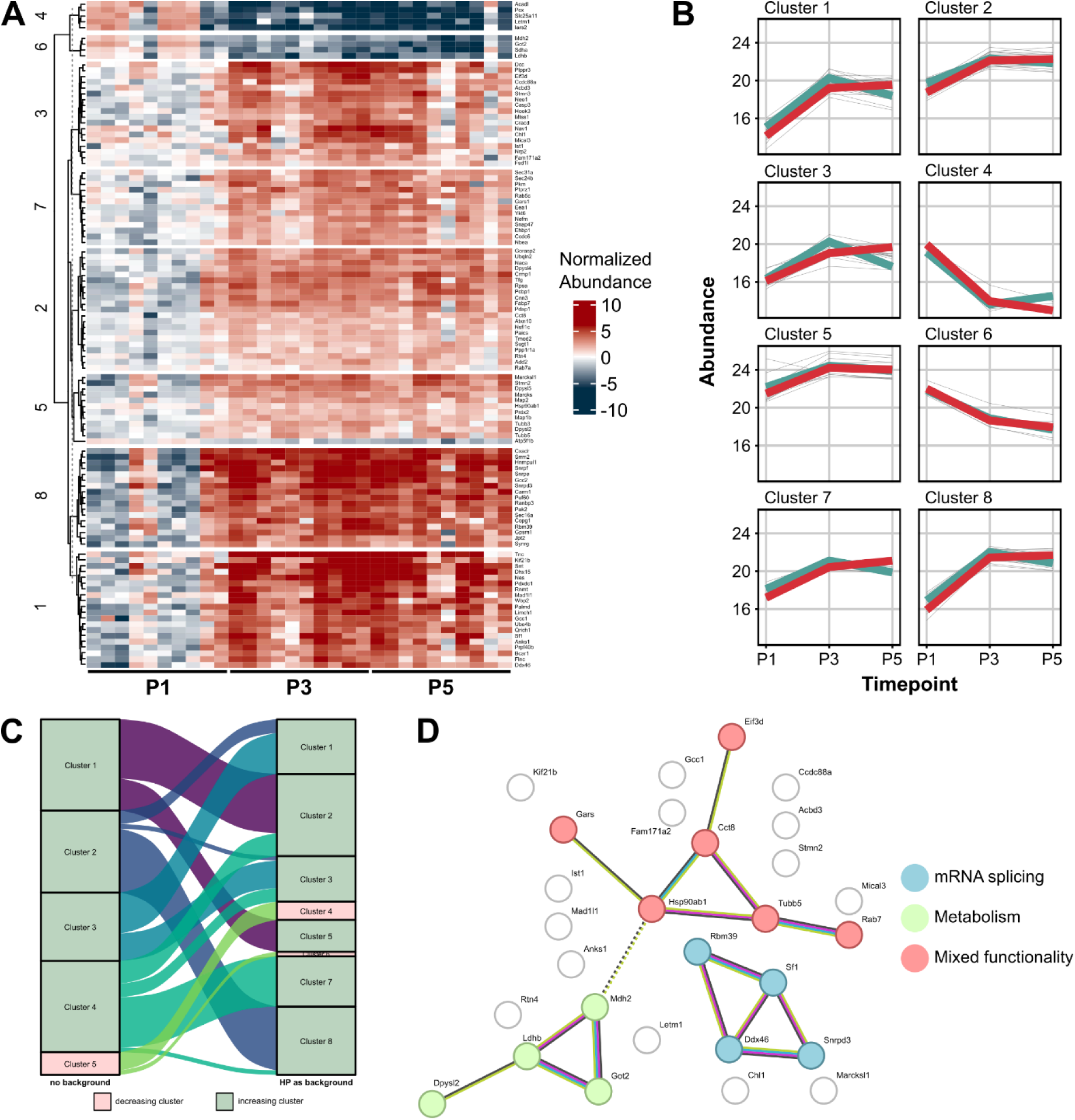
A) Results of a multiple time series analysis with HP as control. The heatmap shows the normalized individual protein abundances and their cluster association. Protein names and their corresponding clusters can be found in the list in STab. 1. B) Average abundances over time identified in A, with the average for HP-GC in teal and the average for DG-GCs in red. Protein individual curves are grey. C) Alluvial plot on clustering consistency between the single time series and multi time series analysis of shared proteins. Clusters with decreasing abundance are colored in pink and clusters with increased abundance in green. D) Results for STRING network analysis of proteins only identified as significantly changed in the multi-time series analysis.

To evaluate if identical proteins were identified in the single- and multi-time series analysis, we compared the list of temporally regulated proteins of both evaluations. We found 28 proteins that were identified only in the analysis of DG-GCs against HP-GCs. The multi-time series analysis shared four proteins with the single-time series for the HP, 74 proteins with the DG, and four proteins with both. The high overlap with the single-time series analysis of DG-GCs was expected, as only a small number of temporary, significantly changed proteins were identified for the HP-GCs themselves, thus, the majority of those proteins could originate from the large group of temporary regulated proteins in DG-GCs. To ensure the consistency in proteins trajectory (e.g. same proteins showing increasing or decreasing trajectories in both analyses), we visualized the clusters in an alluvial plot and confirmed that no protein moved from an increasing cluster to a decreasing cluster and vice versa between the two analyses (Figure 6C).

To evaluate the interaction for the 28 proteins additionally identified in the multi-time series analysis we run a STRING network analysis that resulted in three distinct functional networks (Figure 6D), which are associated with mRNA splicing, metabolism, or mixed functionality around protein folding and stability (HSP90AB1 (Haase & Fitze, 2016) and CCT8 (Kelly et al., 2022)) and neuronal differentiation and dendritic spine formation (TUBB5 (Ngo et al., 2014) and RAB7 (Shikanai et al., 2018)). The only proteins that do not show an increase in abundance in the multi-time series analysis are GOT2, LDHB, MDH2 – all associated with metabolism – and LETM1, a protein important for maintaining mitochondrial morphology (Jiang et al., 2013). In the metabolism network DPYSL2, also known as CRMP2 and a well-known GC marker, was increased in abundance, which can be linked to its function in axon growth and guidance (Marques et al., 2013). Furthermore, DPYSL2 contains a nuclear localization sequence that could be indicative that DPYSL2, or one of its isoforms (Feuer et al., 2023), has another function in GC-nucleus communication.

The comparative analysis of HP-GCs and DG-GCs reveals subregion-specific developmental programs. While most temporal regulated proteins in HP-GCs show peak abundance around P3 (Figure 6E, Cluster1, Cluster 3, Cluster 4, Cluster 7, and Cluster 8), DG-GC proteins exhibit more sustained abundance without a clear P3 peak. Remarkably, six out of eight clusters in the multi-time series analysis show increasing trajectory in protein abundance, which reflects intensive biosynthetic activity characteristic for critical developmental windows. The decrease of metabolic proteins from P1 to P5 may indicate a developmental shift from more general metabolic support to more specialized neuronal functions as proteins associated with metabolic processes decrease in their abundance while at the same time those important for modifying local translation and protein stability increase. This indicates a shift from an energy-producing and consuming phase to one in need of a changed, new subset of proteins and function.

### 6. Proteins associated with neurological disorders

The hippocampus, like other brain regions, relies on the proper establishment of the cellular connections of its circuit. This depends on the accurate pathfinding of the respective GCs responsible for connecting the different subregions within the circuit. Disruption in GC dynamics, cytoskeletal regulation, and axonal guidance can contribute to both neurodevelopmental disorders and in the long term to neurodegenerative diseases (Weerasinghe-Mudiyanselage et al., 2022). We used disease ontology (DO) analysis on the whole dataset (HP-GC and DG-GC proteome) to identify proteins in our dataset that are associated with neurological disorders and neurodegenerative diseases (Table 1). We identified over 400 proteins in our dataset that are associated with 31 different neurological dysfunctions. Approximately 20 of those proteins were identified in our time series analyses and are of special interest, because those proteins might exhibit key functions during the development of the HP and DG. Among them is CASP3, well known for its various roles in several diseases (McIlwain et al., 2013) and the GC marker DPYSL2, that is associated with epilepsy and seemed to be especially important for the development of the DG. The largest overlap with time regulated proteins were found for Alzheimer’s disease, Amyotrophic lateral sclerosis (ALS), autism spectrum disorder, and epilepsy syndrome. Several studied suggested that defects in the axonal RNA translation may contribute to the progression of ALS, highlighting the importance of the RNA function associated proteins in our dataset (Cagnetta et al., 2018; Murakami et al., 2015).

**Table 1:**
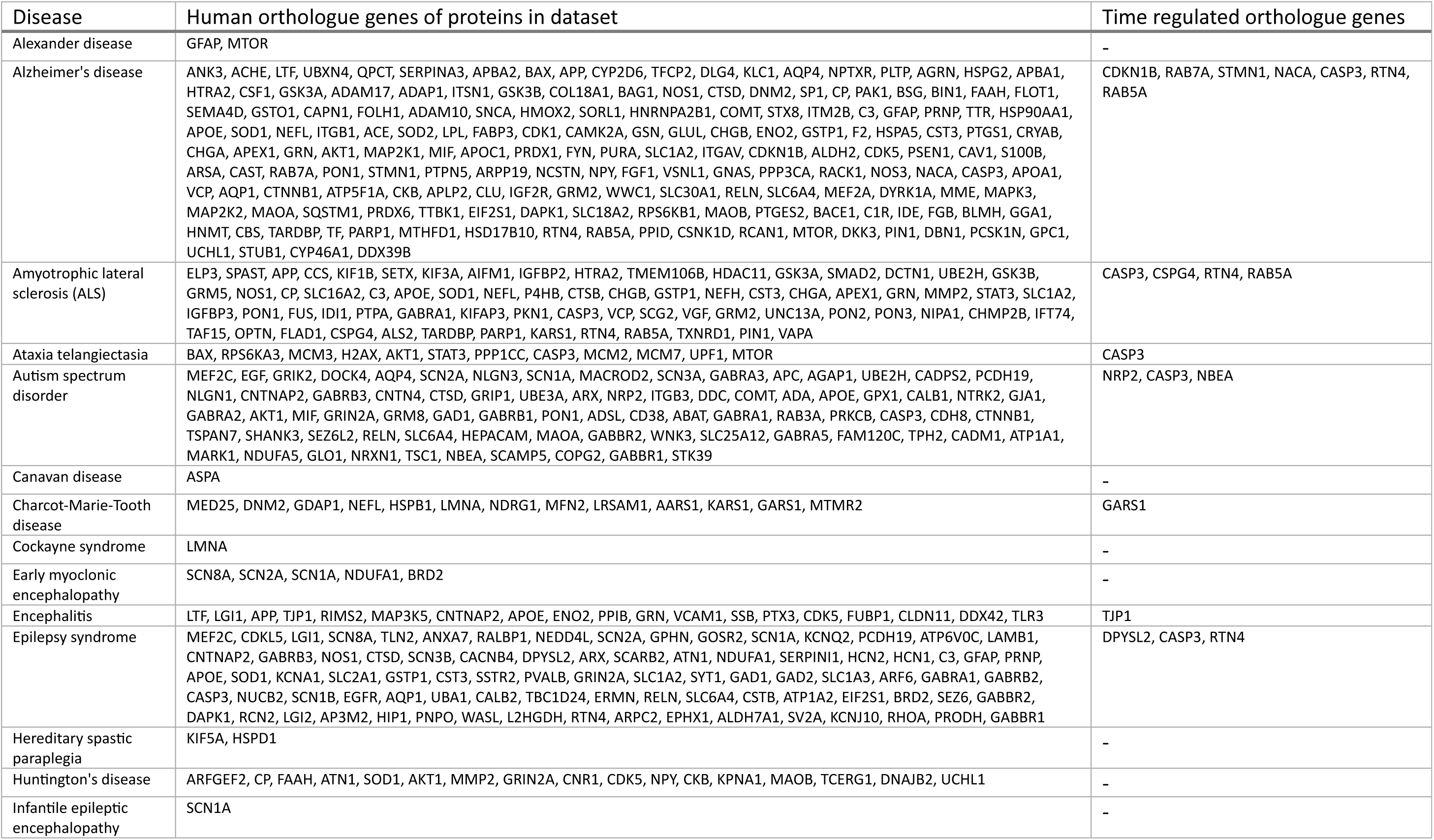
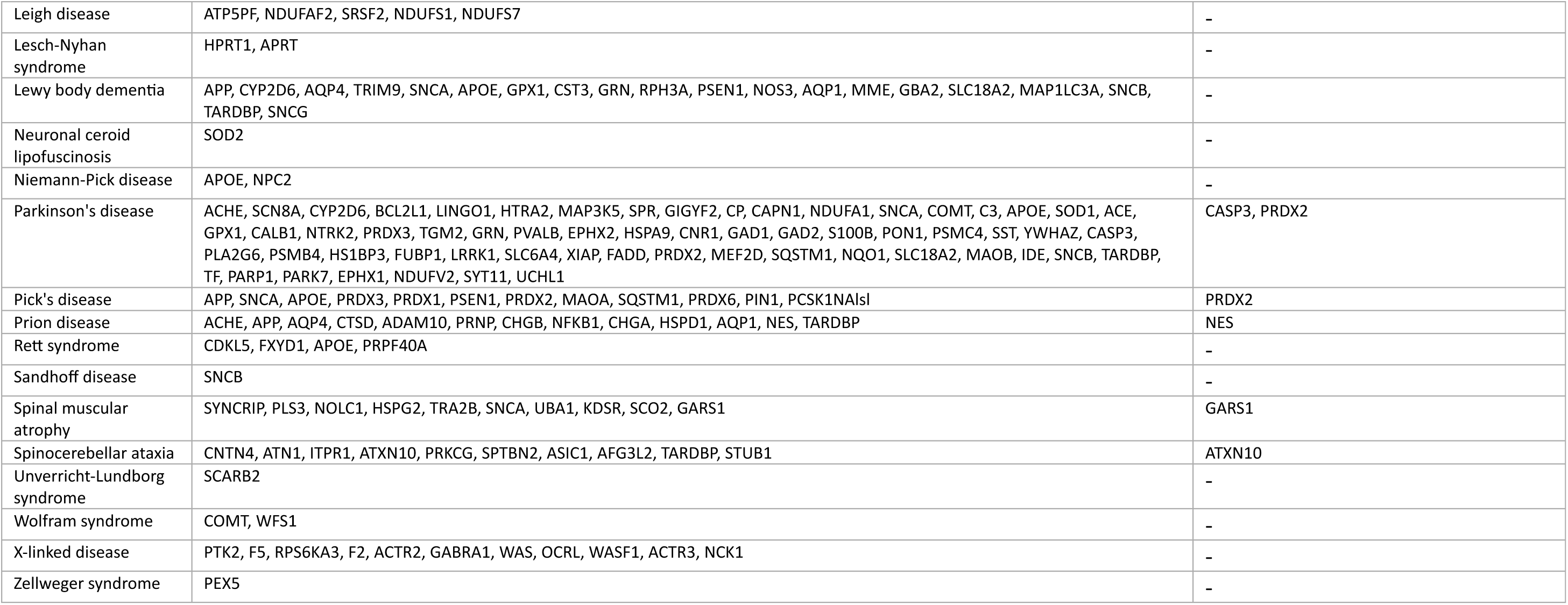
Human orthologues genes represented in our dataset that are associated with neurological and neurodegenerative diseases. A subset of time regulated disease associated orthologous genes is indicated separately. Table depends on disease term enrichment analysis and selection of relevant diseases.

## 4. Discussion

The early postnatal period in rodent development represents a critical window for neural circuit assembly and refinement. Our characterization of hippocampal invasion by entorhinal cortex (EC) layer 2 Reelin-expressing neurons confirmed that the first days after birth are crucial for circuit development. By postnatal days 5 (P5), many connections characteristic for the mature entorhinal-hippocampal circuit have been established. P1, P3, and P5 correspond to particularly dynamic phases of entorhinal-hippocampal circuit development with distinct characteristics. At P1, the basic architecture of the hippocampal formation has been established, though many connections remain immature and not fully in place. Granule cell neurogenesis in the dentate gyrus (DG) continues at high rates during this period, with peak neurogenesis occurring around P5-P7 (Bayer, 1980; Schlessinger et al., 1975). P3 represents a critical transitional period when many developmental processes accelerate. During this phase, topographic projections from the EC undergo refinement, and inappropriate connections begin to be eliminated through activity-dependent mechanisms (Amaral & Dent, 1981; Supèr & Soriano, 1994). Additionally, the laminar termination patterns of entorhinal projections in the DG become more organized and local inhibitory circuits develop rapidly (Amaral & Dent, 1981; Pleasure et al., 2000). By P5, coordinated network activity patterns emerge, which are thought to drive circuit maturation and functional development (Ben-Ari et al., 1989; Khazipov & Luhmann, 2006). This period represents a critical time for experience-dependent plasticity mechanisms that shape the final circuit organization.

In our study we focused on the first five postnatal days and extracted GCs from the HP and the DG to study their proteomic composition. We identified over 5000 proteins in total for the different GC preparations and with each timepoint the HP-GCs and DG-GCs were characterized by a distinct protein subset (Figure 3B and Figure 4B). Interestingly, the timepoint specific proteins covered very similar biological processes in HP and DG at P1, but with increasing age those processes diverge, clearly demonstrating that the DG develops on a slower timeline than the HP overall (Figure 5C). This temporal offset was particularly evident in the expression of proteins involved in *RNA spicing, via spliceosome*. Proteins associated with spliceosome function appeared at earlier timepoints in HP-GCs but at later timepoints in DG-GCs. The prominence of *Ribosomes* and *Spliceosomes* among the Top20 KEGG pathway terms for proteins shared across all timepoints underscores two critical aspects of GC biology: first, it highlights the importance of local protein synthesis for growth cone function; second, it demonstrates the capacity for rapid proteome modulation through alternative splicing, which enables fast responses to local environmental cues and guidance signals (Cagnetta et al., 2018; Estrada-Bernal et al., 2012; Shigeoka et al., 2016).

The overall shift in biological processes across developmental stages indicates that GCs mature from a mobile, exploratory state toward one focused on synaptic maturation and stable connection formation. Notably, P1 was majorly characterized by metabolic functions, particularly fatty acid metabolism. Among the temporally regulated proteins in DG-GCs we identified the lipid-interacting protein FABP7, which modulates the availability of fatty acids that serve as precursors for membrane synthesis, indicating its involvement in coordinating lipid metabolism with neuronal maturation (Young et al., 2013). The importance of fatty acid metabolism for GCs was also demonstrated in previous studies (Chauhan et al., 2020). Given that FABP7 deficiency has been associated with impaired hippocampal neurogenesis and altered behavioral outcomes (Young et al., 2013), disruptions in FABP7-mediated lipid metabolism during these critical early postnatal days could have lasting consequences for circuit development and function.

Our temporal trajectory analysis for HP-GCs revealed fifteen proteins that show a significant difference in their expression over time. While this number may seem surprisingly low, it likely reflects the cellular heterogeneity within our samples. The HP-GC preparations contained GCs of multiple cell types extending axons toward their respective targets, including projection neurons from EC layers 2 and 3 granule cells from the DG (Mossy fibers), and projecting neurons from the CA regions (e.g. Schaffer collaterals). Thus, the fifteen temporally regulated proteins likely represent those with fundamental importance for overall HP development. Notably among these, we identified two GC markers, DCC and SEC31A. DCC has been shown to cover multiple functions, particularly in axon guidance (Powell et al., 2008; Ren et al., 2007). In the context of high translational activity observed in our dataset, DCC’s ability to bind and regulate local translation is especially significant (Koppers et al., 2019; Koppers & Holt, 2022). Importantly, DCC was also identified as temporally regulated in DG-GCs, indicating its general importance across all HP subregions.

Another interesting GC marker in our dataset is DPSYL2 (also known as CRMP2), which exhibits manifold functions ranging from axon guidance and cytoskeletal interactions to endocytosis and neurotransmission (Feuer et al., 2023). A less-characterized function of DPSYL2 relates to its nuclear localization sequence (NLS). NLS are short signal peptide sequences that mediate the transport of proteins into the nucleus (Lu et al., 2021), suggesting a potential role in transcriptional regulation and hinting at direct communication between GC and nucleus. Other NLS-containing proteins in our dataset include HNRNPH2 and FABP7. HNRNPH2, a member of the heterogeneous ribonucleoprotein family, participated in various aspects of RNA metabolism, such as RNA processing, alternative splicing, and mRNA trafficking. It uses its nuclear localization to access pre-mRNA, before being shuttled out of the nucleus (Korff et al., 2023). FABP7 mediates lipid-dependent transcriptional regulation after binding fatty acids (reviewed in Needham et al., 2022). The presence of temporally regulated, NLS-containing proteins in our dataset raises important questions about the interaction between GCs and their nucleus, and the functional consequences of this direct communication for GC maturation. This observation suggests a dual capacity for certain proteins that execute local functions while simultaneously communicating extracellular cues to the nucleus for transcriptional adaptation. Such bidirectional communication may be particularly critical during the developmental window we studied, when GCs must rapidly adapt to changing guidance cues in the HP while coordinating gene expression programs responsible for circuit maturation. This concept is exemplified in a recent study where the role of the transcription factor myocyte enhancer factor 2-c (MEF2C) in postnatal developing neurons was described (Sudarsanam et al., 2025). Knockdown of MEF2C lead to an increased expression of Ephrin A5 (EFNA5), a protein responsible for repulsing guidance cue in normally developing animals. The overexpression in turn resulted in reduced collateral targeting (Sudarsanam et al., 2025). This study elegantly highlighted the interplay between nuclear transcription and axon guidance mediated by the same proteins within individual neurons. Interestingly, we identified MEF2C as one of the temporally regulated proteins for DG-GCs and it will be fascinating to investigate in the future the role of this transcription factor in GCs.

Furthermore, our dataset contains over 400 proteins associated with neurological and neurodegenerative diseases, with twenty showing temporal regulation. Disruptions in GC function have been implicated in various neurodevelopmental disorders, including autism spectrum disorders, amyotrophic lateral sclerosis, and epilepsy (Penzes et al., 2011; Gualdoni et al., 2013). This knowledge is particularly relevant given the growing recognition that many neurological and psychiatric disorders have their origins in developmental disruptions that occur during critical periods of circuit formation (Parenti et al., 2020).

This study has several limitations that should be considered when interpreting our findings. Our proteomic samples contained GCs from multiple cell types, and consequently, we cannot definitively attribute individual proteins and their functions to specific cell types within the HP. While comparing HP-GCs with DG-GCs allows us to speculate about DG-specific proteins, we cannot determine whether these proteins function primarily in, for example, extending mossy fibers or invading layer 2 fibers. To allow more cell specific conclusions on proteins and their function in their corresponding GCs, more refined extractions, e.g. based on introduced fluorochromes, could be applied (Poulopoulos et al., 2019). Additionally, we did not consider the sex of the animals during sample preparation. Both, male and female animals were randomly used for HP-GC and DG-GC preparations without ensuring balanced representation between the sexes within the two sample cohorts. Therefore, we cannot draw conclusions about potential sex-dependent differences in hippocampal formation development.

In summary, our study provides a comprehensive characterization of the GC proteome originating from the whole HP and the DG during early postnatal development. Analysis of timepoint-specific and temporally regulated proteins revealed a clear shift in biological processes within GCs over time. For HP-GCs, P3 emerged as a particularly critical timepoint, with most proteins peaking in expression during this period. In contrast, DG-GCs exhibited a more prolonged developmental trajectory compared to HP-GCs, consistent with the known differences in maturation timelines between these regions. Our findings underscore the dynamic nature of the GC proteome during early postnatal hippocampal development and provide a foundation for understanding how local protein composition guides circuit assembly. The identification of temporally regulated disease-associated proteins highlights the potential importance of these developmental windows for understanding the origins of neurological disorders and suggests that this dataset may serve as a valuable resource for future studies investigating developmental mechanisms of disease pathogenesis.

## 5. Methods

### 1. Animals

Mice belonging to the Odz3-tta mouse line (originally published as MEC-13-53A in (Blankvoort et al., 2018)) were bred with a tetO-GCamp6 reporter mouse line (Blankvoort et al., 2018). Mice positive for both transgenes originating from this cross were used for experiments in Figure 1. Growth cones isolation and control samples were done from wild type mice (C57BL/6JBomTac, Taconic). All animals were housed in enriched environment cages in a reversed 12-hour light/dark cycle with humidity and temperature-controlled housing rooms. Food and water were provided ad libitum. Both male and female mice were used for experiments in this study. All experiments were conducted in compliance with protocols approved by the Norwegian Food Safety Authorities and European Directive 2010/63/EU (FOTS ID 24847; 31016).

### 2. Euthanasia

For euthanasia, all animals were first anesthetized with isoflurane before being euthanized. Pups were euthanized by decapitation and adult animals with a lethal intraperitoneal injection of pentobarbital (100 mg/kg).

### 3. Immunohistochemistry

After euthanasia, animals were transcardially perfused using 5-6 mL of phosphate-buffered saline (PBS) followed by 8-9 mL of 4% (w/v) paraformaldehyde (PFA) in PBS at a flow rate of 3.8-4.0 mL/min. Post-perfusion, brains were extracted and post-fixated in PFA for a minimum of 4 h, followed by 24 h in PBS. The brains were then sequentially incubated for 24 h each in 15% (w/v) and 30% (w/v) sucrose in PBS. Subsequently, the brains were embedded in optimal cutting temperature (OCT) mounting medium and rapidly frozen in a 70% (v/v) ethanol bath. Horizontal brain sections were prepared using a cryostat (CryoStar NX70, Thermo Scientific) at a thickness of 20 µm, directly mounted on adhesion slides (Superfrost Plus Adhesion Microscope Slides, Epredia), and stored at −20°C until further usage. For immunohistochemistry, the adhesion slides were thawed and framed using a PAP Pen (5 mm tip, Merck) before being rinsed in PBS for 10 min. Blocking was performed for 30 min in blocking solution (1X PBS, 0.1% (v/v) Triton-X, 10% (v/v) donkey serum). The slides were then incubated overnight at 4°C with the following primary antibodies: NeuN (Guinea Pig, anti-NeuN, Sigma Millipore #ABN90P, 1:1000), GCamP6 (Chicken, anti-GFP, Abcam #AB13970, 1:1000), and Stx7 (Rabbit, anti-Stx7, Synaptic Systems #110072, 1:200). Antibodies were diluted in incubation buffer (1X PBS, 0.1% (v/v) Triton-X, 1% (v/v) donkey serum). The following day, slides were rinsed 6 x 5 min in PBS and incubated for 2 h with the following secondary antibodies: donkey anti-Guinea pig 405 (Jackson Immuno Research Lab #706-475-148, 1:250), donkey anti-Chicken 568 (Invitrogen #A78950, 1:500), and donkey anti-Rabbit 647 (Invitrogen #A31573, 1:500), diluted in incubation buffer. After rinsing the slides 6 x 5 min in PBS, they were coverslipped with Flouromount-G (Invitrogen) and sealed with standard transparent nail polish.

### 4. Confocal Imaging and growth cones count

The HP and EC were imaged in 10x (Plan-Apochromat 10x/0.45 M27) and 40x magnification (Plan-Apochromat 40x/1.4 oil DIC M27) using the LSM 880 confocal microscope (Zeiss). Tiled *z*-stacks were acquired, and a 3D image was reconstructed from the *z*-stack layers. In total, three horizontal sections from each of three animals (n = 9) were chosen for imaging. Sections used for counting were a minimum of 150 µm apart. All images were processed in Zen software (blue edition, version 2.6.76, Carl Zeiss Microcopy, 2018) and further deconvoluted using Huygens Essential 23.10 software (Scientific Volume Imaging). Deconvoluted confocal imaging files containing the hippocampus were uploaded to Neurolucida 360 (Micro Bright Field Bioscience) for analysis. To analyze the fluorescent intensity of invading EC Reelin-expressing layer 2 fibers and the amount of growth cones in the hippocampus, contours delineating the stratum lacunosum-moleculare of CA3 (CA3 SLM), and the molecular layer (ML) of the inner blade and of the outer blade of the Dentate Gyrus were created. Growth cones that were both positive for GCamP6 and Stx7 were labelled with markers and counted. Since non-consecutive slices were analyzed, overcounting the z-axis was not relevant, and no correction was applied. For the quantification of the number of count markers and the fluorescent signal within the delineated areas, contours and counts were exported as xml files and quantified using a customized MatLab (R2023a version, Mathworks) script. Briefly, the cartesian coordinates for contours and counts were extracted from the xml format Neurolucida 360 files and transformed into a Boolean mask fitted to match the size to the original reference picture on which the delineations were drawn. The masks were used to extract the fluorescence intensity values of the topologically corresponding voxels from the reference confocal image. A squared region with a standardized size (400×400 pixels) located in the weighted midpoint (centroid) of the Dentate Gyrus was chosen as the background region. A threshold value was set as being the sum of the average value plus three times the standard deviation of fluorescence intensity of the background region for each image. The number of voxels within each contour (CA3 SLM, inner LM, and outer LM) in which the fluorescence intensity exceeds the threshold value were counted (SFig. 1B). Voxel counts were then normalized to the total number of voxels for the corresponding delineated area. Outliers within the boxplots were identified based on the quartile methods and labeled with asterisk.

### 5. Hippocampus and Dentate Gyrus Microdissections

Animals at P1, P3, and P5 were euthanized and decapitated. The brains were promptly extracted from the skull, and the regions of interest were dissected on ice under Stereo Microscope SZ16 (Olympic Life Sciences) in ice-cold growth cone homogenization (GCHO) buffer consisting of 4 mM Hepes, 0.32 M Sucrose (pH 7.4) supplemented with 100 µL of 0.5 M EDTA stock and 100 µL Halt protease inhibitor cocktail per 10 mL (Thermo scientific). First, the cerebellum was removed and the hemispheres separated by a midline incision along the longitudinal fissure. With the medial side facing up, the white matter was excised, exposing the hippocampus. The hippocampus was carefully flipped out of the cortex using a small paint brush or forceps. The fibers connecting the cortex to the hippocampus were severed. For the DG samples, the HP was subsequently carefully flipped the medial side up, and the DG was isolated by gently separating it from the CA, using as reference the visual gap between the DG and CA regions, along the septotemporal axis. The HP and DG samples were either frozen for RNA extraction the following day or immediately immersed in ice-cold GCHO buffer and stored on ice for growth cone preparation the same day.

### 6. qRT-PCR

RNA was extracted from microdissected DG and HP P4 tissues using the NucleoSpin RNA/Protein kit (Machinery-Nagel) following the manufacturers protocol. Reverse transcription of 100 ng of RNA into cDNA was performed using the Eurogentec RT-RTCK-05 Reverse Transcriptase Core kit 500. qRT-PCR was conducted in 20 μL reaction containing 12 μL SensIFAST SYBR mix and 8 μL cDNA (1:10 diluted RT reaction), using StepOne Real-Time PCR system (Applied Biosystems). Cycler conditions included initial denaturation at 95°C for 2 min, followed by 40 cycles of denaturation at 94°C for 5 sec, annealing at 60°C for 10 sec, and extension at 70°C for 20 sec. Cт values were averaged from technical duplicates, excluding values above 35. Relative gene expression was calculated using the 2(−ΔΔCт)-method (Schmittgen & Livak, 2008), normalized to *Gus* (D. Xu et al., 2018). Primer specificity for housekeeping gene *Gus* was confirmed through NCBI *BLAST searches specific to Mus musculus*. DG and CA specific target genes are *Neurod1, Neurod4*, *Dusp14*, and *Meis2*, *Neurod6*, and *Tyro3*, respectively. Primers, listed in Table 2, were designed using NCBI Primer-Blast with a target product length of 70 – 150 bp. Statistical significance of mRNA levels between CA and DG were detected by independent two tailed Student’s t-test.

**Table 2:**
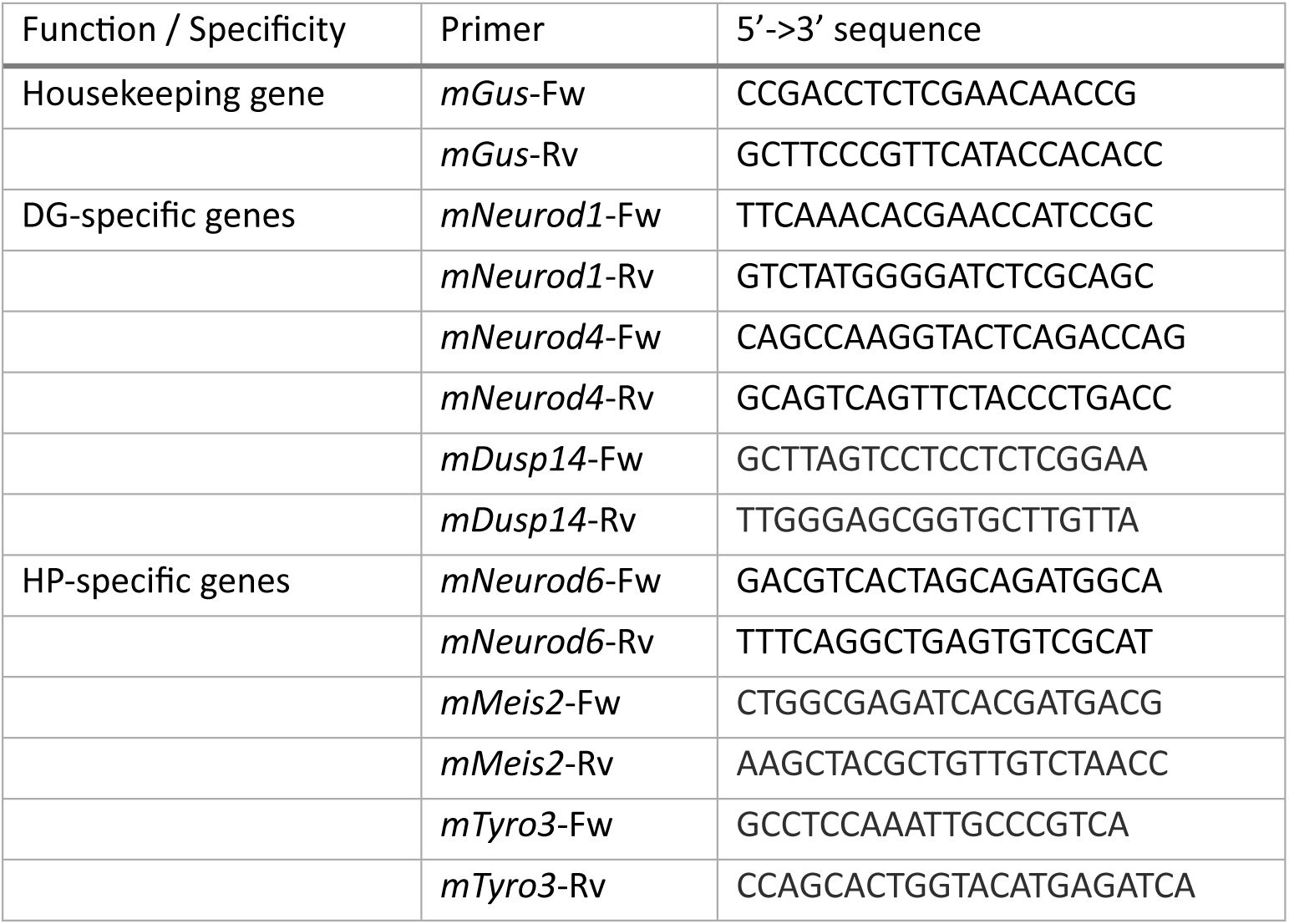
5’->3’ sequences and function or tissue specificity of primers used for qRT-PCR. Fw stands for forward and Rv for reverse.

### 7. Growth Cone Preparation

The growth cones were prepared as detailed in (Pfenninger et al., 1983). Briefly, the microdissected tissue was homogenized in GCHO buffer using glass-Teflon homogenizer coupled with Hei-TORQUE overhead stirrer (Heidolph) with thirteen slow and even strokes at 900 rpm. The homogenate was transferred to a reaction tube and centrifuged for 15 min at 1,700 x *g*, at 4°C (Eppendorf, Refrigerated Microcentrifuge 5426). The post-nuclear homogenate (PNH) was separated from the pellet and subsequently diluted with GCHO buffer to a final volume of 1.7 mL. A discontinuous sucrose gradient was layered in ultra-clear centrifugation tubes (5.6 mL, Beckman No: 344075) with three layers: a bottom cushion (4 mM Hepes, 2.5 M sucrose, pH 7.4), a middle layer (4 mM Hepes, 0.83 M sucrose, pH 7.4), and a top layer of the diluted PNH, all with a volume of 1.7 mL. The tube necks were sealed, and the gradient was centrifuged at 250 000 x *g* for 50 min at 4°C using a Sorvall Discovery 100SE (Thermo Scientific) or Optima XE-90 (Beckman Coulter) equipped with a VTi50 Rotor (Beckman Coulter). Following centrifugation, growth cone particles (F1 fraction) were recovered from the interface between the middle and top sucrose layer using an 18-gauge needle and a 3 mL syringe. The F1 fraction was either transferred onto a polycarbonate membrane filter (Millipore, No: VCT02500, pore size 0.1 µm), filtered with Rotavac Valve Control vacuum pump (Heidolph, No: 591-00130-00-2), and stored at −80°C until analysis by LC-MS/MS or diluted in PBS for subsequent Western blot analysis.

### 8. Western blot

Growth cone preparation was validated by western blot. F1 fractions were diluted 1:5 in PBS and pelleted at 12,000 x g, 5 min centrifugation. Pellets were resuspended in PBS and mixed with 4x Lämmlie buffer β-mercaptoethanol and incubated at 95°C for 5 min. Synaptosome and total hippocampal protein extract control samples were prepared following the synaptosome preparation by Dunkley (Dunkley et al., 2008) and using RIPA buffer (Thermo Fisher Scientific) extraction according to the manufacture’s protocol. Both types of control samples were denatured with Lämmlie buffer as described above. Protein concentrations were detected using the G-Biosciences Compatible Lowry assay. 1 µg of each sample was separated via SDS-PAGE on 8-16% Mini-PROTEAN TGX Stain-Free gels (Bio-Rad) under the manufacturer’s 200 V standard protocol. Proteins were transferred to PVDF membranes, blocked with TBS-T (1x TBS, 0.1 (v/v) % Tween-20) containing 5 (w/v) % skim milk powder for 1 h at room temperature, and probed overnight at 4°C with primary antibodies diluted in TBS-T with 1 (w/v) % skim milk powder. The primary antibodies used were: Rabbit anti-Growth associated protein-43 (1:1000, Merck #AB5220), Rabbit anti-Synaptophysin antibody (1:2500, Abcam #14692), Rabbit anti-Map2 polyclonal antibody (1:1000, Proteintech #17490-1-AP), Mouse anti-PSD95 antibody [K28/43] synaptic marker (1:2500, Abcam #192757), and Mouse β-Actin loading control monoclonal antibody (BA3R) (1:1000, Invitrogen #MA5-15739). After washing (3 × 10 min in TBS-T), the membrane was incubated for 1.5 h at room temperature with the following HRP-conjugated secondary antibodies: Goat anti-Rabbit IgG (H+L) (1:2500, Promega #W4011) and Rabbit anti-Mouse IgG (H+L) (1:5000, Invitrogen #31450), diluted in TBS-T with 1% skim milk powder. The membrane was washed (3 × 10 min in TBS-T), incubated in ECL solutions (Clarity Western Peroxide Reagent and Enhancer Reagent, Bio-Rad) for 2 min, and chemiluminescence was detected using the ChemiDoc XRS+ system and ImageLab software (Bio-Rad).

### 9. LC-MS/MS

Mass spectrometry sample analysis was performed by the Proteomics and Modomics Experimental Core Facility (PROMEC, NTNU). Tryptic samples were prepared from polycarbonate filters by first recovering the GC in 600 µl 1% SDC, 100 mM Tris-HCl pH 8.5, 10 mM TCEP, 40 mM CAA and sonicated for 30 min 70°C. Afterwards, 600 µl 0.1 M Ammonium bicarbonate, 0.5 µg trypsin were added and the samples digested over-night at 37°C. Peptides were desalted using C18 spin columns, dried in a speedvac centrifuge and resuspended in 0.1% Formic acid before MS analysis. LC-MS/MS were performed on a timsTOF Pro (Bruker Daltonics) connected to a nanoElute (Bruker Daltonics) HPLC. The peptide solution was injected, and separation was done using a Pepsep 25 (150 µm x 25 cm) column with running buffers A (0.1% Formic acid) and B (0.1% Formic acid in Acetonitrile) with a gradient from 0% B to 40%B for 75 min. The timsTof instrument was operated in DDA PASEF mode with 10 PASEF scans per acquisition cycle and accumulation and ramp times of 100 ms each. The ‘target value’ was set to 20,000 and a dynamic exclusion was activated and set to 0.4 min. The quadrupole isolation width was set to 2 Th for m/z < 700 and 3 Th for m/z > 800.

### 10. Proteomic data analysis

TimsTOF DDA raw data files were interpreted using *MaxQuant* software v. 2.6.1.0 (Cox & Mann, 2008). Proteins were identified with one unique peptide and a false discovery rate of 0.01 using the mouse proteome including isoforms from Uniprot (downloaded: 14.03.2024) as a reference dataset. Analysis parameters were Trypsin/P as a digestion enzyme; a maximum of two missed cleavages were allowed; six amino acids were used as minimum peptide length; oxidation (M), acetyl (protein N-term), and deamidation (NQ) were set as variable modifications. Label-free quantification (LFQ) was performed with match between runs enabled using MaxQuant standard settings. *PTXQC* R package (Bielow et al., 2016) was used to quality control results generated by MaxQuant. All further downstream analyses were conducted using a custom R script (R v. 4.2.3, script available from XXXXX). All proteins labelled as *potential contaminants*, *only identified by site*, and *reverse* were removed from the dataset. Proteins were considered as identified for a timepoint and tissue type if three out of five biological replicates showed an LFQ intensity value. GO term analyses were performed using *ClusterProfiler* v. 4.16.0 (S. Xu et al., 2024). GO term results were manually curated and redundant terms triggered by identical proteins removed. KEGG pathway analyses for mouse protein terms were performed using the online platform *ShinyGO 0.85* (Ge et al., 2020). For single and multi-series time course analyses the package *maSigPro* v. 1.81.0 (Conesa & Nueda, 2025) was used with a quadradic regression model (degree = 2) and the parameters Q = 0.05, Benjamini-Hochberg correction for multiple testing adjustment, and a minimum of four observations for single series time course analysis using P1 as reference and the parameters Q = 0.05, Benjamini-Hochberg correction for multiple testing adjustment, and a minimum of six observations for the multi-series time course analysis with the hippocampal data as reference dataset. Human orthologues genes were identified using *orthogene* v. 1.14.01 (Schilder & Skene, 2022). Disease terms were identified by using *DOSE* v. 4.2 (Yu et al., 2015). All plots were generated in R using the packages *ggplot2* v. 3.5.2 (Wickham, 2016), *ggVennDiagram* v. 1.5.2 (Gao & Dusa, 2025), *alluvial* v. 0.1-2 (Bojanowski & Edwards, 2016), and *ComplexHeatmap* v. 2.24.1 (Gu et al., 2016). Raw data is available at the ProteomeXchange consortium (http://proteomecentral.proteomexchange.org) submitted via the PRIDE partner repository as PX Partial with the dataset identifier XXXX (Deutsch et al., 2017; Perez-Riverol et al., 2022).

## Author Contributions

Conceptualization, M.K. and G.Q.; supervision, M.K. and G.Q.; experiment design, M.K. and G.Q.; data collection, M.K., K.A.K., and L.S.S.; data evaluation, M.K., K.A.K., and P.J.B.G.; visualization, M.K. and P.J.B.G.; writing, original draft, M.K.; writing, review, and editing, M.K., K.A.K., P.J.B.G, L.S.S., and G.Q.; funding acquisition: G.Q.; discussion, comments, all authors.

## Acknowledgement

We thank Bianca Ana Zaharia and Julie Six for outstanding technical assistance and the whole animal technician team at Kavli institute for Systems Neuroscience. LC-MS/MS sample preparation and analysis were performed by the Proteomics and Modomics Experimental Core Facility (PROMEC) at NTNU. PROMEC is funded by NTNU and the Central Norway Regional Health Authority and a member of the National Network of Advanced Proteomics Infrastructure (NAPI), which is funded by the RCN INFRASTRUKTUR-program (295910). The work was supported by a Research Council of Norway (RCN) FRIPRO grant to G.Q. (grant number 324305), the Trond Mohn Foundation (Mohn Research Center of the Brain, grant number 2021TMT04, to G.Q.), an RCN Centre of Excellence grant (Centre for Algorithms in the Cortex, grant number 332640; Centre of Neural Computation, grant number 223262; to G.Q.), and the Kavli Foundation (to G.Q.). The experiments were performed at the NORBRAIN Facility, Norwegian University of Science and Technology (NTNU) (grant number 295721).

